# Multi-layered apoplastic barrier underlying the ability of Na^+^ exclusion in *Vigna marina*

**DOI:** 10.1101/2024.04.23.590519

**Authors:** Fanmiao Wang, Keitaro Tanoi, Takaki Yamauchi, Ken Naito

## Abstract

- Salt tolerance is important to tackle problems of soil salinization and ground water depletion. However, developing salt tolerant crops is facing difficulties due to limited potential in model plants and crop species. Thus, it is important to elucidate how coastal species, such as *Vigna marina*, adapt to saline environments.
- By comparative transcriptome and histological analyses, this study elucidated one important aspect of how *Vigna marina* achieves salt exclusion and extraordinary salt tolerance.
- Under salt stress, genes involved in casparian strip formation were specifically upregulated in JP247202 (*V. marina*). JP247202 reinforced apoplastic barrier with thick lignification in multiple layers of cells around endodermis. Also, disruption of lignification led to a dramatic increase of shoot Na^+^ accumulation and salt lesion in JP247202. Interestingly, despite the salt-induced apoplastic barrier, JP247202 maintained transport of essential ions including K^+^, Mg^2+^, and Ca^2+^.
- Our results revealed that lignification of multi-layered cells around endodermis was an important apoplastic barrier to the transport of Na^+^ to shoots in JP247202, while it did not restrict the transport of K^+^, Mg^2+^, and Ca^2+^. This feature, together with the ability of Na^+^ excretion by SOS1, enables *V. marina* to thrive in marine beaches.

## Introduction

Wild plants have great potential for stress tolerance. Wild relatives of crop species preserve the original genetic variation, which is a prerequisite for crop breeding and has been lost in cultivated species during the process of domestication and breeding (Hufford *et al*., 2019; Gasparini *et al*., 2021). For example, more than half of the genetic variation has been lost in cultivated soybean (Hyten *et al*., 2006; Zhou *et al*., 2015) and genetic diversity has been significantly reduced in cultivated rice (Xu *et al*., 2012). Moreover, breeding efforts mainly aim for high yield, harvesting, and edibility (Konishi *et al*., 2006; Hua *et al*., 2015; Zhang *et al*., 2018), rendering modern varieties vulnerable to biotic and abiotic stresses (Rosenthal and Dirzo, 1997; Gilliham *et al*., 2017). For example, cultivated wheat and barley are less adaptive to drought and salt compared to their wild progenitors (Nevo and Chen, 2010) and the domesticated tomato is more sensitive to salt stress than its wild relatives (Wang Z. *et al*., 2020). Stress tolerance of wild plants could be introduced to susceptible modern crops through conventional breeding or transgenesis. Therefore, it is necessary to understand mechanisms of stress tolerance and underlying genetic factors in wild plants.

Wild species in the genus *Vigna* thrive in harsher environments thus they must have different or reinforced strategies for adaptation. In fact, compared to crop species in the genus *Vigna*, including azuki bean, cowpea and etc., wild species have higher salt tolerance (Yoshida *et al*., 2020). Among them, *Vigna marina*, growing by the sea, is a highly salt-tolerant species that can survive in 300 mM of NaCl for at least four weeks (Yoshida *et al*., 2020). This is remarkable given that few of the current crops can tolerate 120 mM of NaCl (Kotula *et al*., 2020). The most notable feature of *V. marina* is the suppression of salt accumulation in shoots (Noda *et al*., 2022), maintains high transpiration and photosynthesis (Yoshida *et al*., 2020) and water uptake under salt stress (Wang *et al*., 2023). *Vigna luteola* is another wild species in the genus *Vigna*. Different from *V. marina*, not all accessions of *V. luteola* are growing at the seaside thus salt tolerance varies among accessions. Compared to *V. marina, V. luteola* accumulate more salt in shoots, while patterns of salt accumulation in stems and each leaf differs between tolerant and sensitive accessions (Noda *et al*., 2022; Wang *et al*., 2023; Noda *et al*., 2024). The genetic mechanism of salt tolerance remains elusive until a recent study revealed that *V. marina* has elevated activity of *SOS* pathway thus greater ability to excrete Na^+^ out of roots (Noda *et al*., 2024).

However, given that the QTL containing *SOS1* locus explains only 10% of phenotypic variance (Noda *et al*., 2024), this study aims to seek for other mechanisms of the salt tolerance in *V. marina*. We show that the expansion of lignification to multi-layered cells around endodermis is one of the important components for the excellent salt exclusion of *V. marina*. Disruption of lignification led to an increased Na^+^ accumulation in shoots and salt damage of *V. marina*. This knowledge learned from the outstanding salt-tolerant wild plant will bring new insights into developing salt-tolerant crops.

## Materials and methods

### Plant materials

Three accessions of the genus *Vigna* were used in this study: JP247202 (*Vigna marina*), JP233389 (*V. luteola*) and JP235855 (*V. luteola*). Seeds were obtained from the Research Center of Genetic Resources, National Agriculture and Food Research Organization (NARO), Japan (NARO Genebank: https://www.gene.affrc.go.jp/index_en.php).

### Plant growth conditions

Seeds were sterilized with 70 % ethanol and 5 % sodium hypochlorite for five minutes each and rinsed thoroughly with tap water. Sterilized seeds were germinated on Seramis granules (Effem GmbH, Verden, Germany) for one week after scratching on the seed coat to ensure water absorption. Germinated seedlings were transplanted in hydroponic culture containing 1× Otsuka house No. 1 and 1× Otsuka house No. 2) (Otsuka Chemical Co., Osaka, Japan: N, P, K, Ca, and Mg = 18.6, 5.1, 8.6, 8.2 and 3.0 mEq L^-1^, respectively). Plants were grown in a growth chamber at 28 °C with light from 6:00 am–8:00 pm (14 h) and in a dark from 8:00 pm to 6:00 am (10 h). After two weeks, 200 mM of NaCl was treated with a renewed hydroponic culture. Plant growth was investigated daily.

### Retrieving and processing of RNA sequences

Root RNA-seq data of JP247202, JP233389 and JP235855 with BioProject ID of PRJNA1080052 was retrieved from NCBI for a reanalysis. Briefly, this data was collected from three biological replicates in control and salt treatment of 200 mM of NaCl at 24 hours (day 1 (D1)), 36 hours (night 1 (N1)), 48 hours (day 2 (D2)) and 60 hours (night 2 (N2)) (Suppl. Table 1). RNA-seq reads were quantified using the quasi-mapping-based mode of Salmon (Patro *et al*., 2017) with indexing of a de-novo assembly primary transcript of *Vigna marina*. The resulting raw counts were Trimmed mean of M values (TMM) -normalized using EdgeR (Robinson *et al*., 2010) for downstream analyses.

### Analyses of differentially expressed genes (DEGs) and gene ontology (GO) term enrichment

DEGs between control and salt at each time point were determined using EdgeR with a threshold of a false discovery rate ˂ 0.01 and an absolute log2 fold change > 1. DEGs that were shared between time points (D2 and N2 for JP247202; D1, N1, D2, and N2 for JP233389 and JP235855) were defined as common DEGs.

Common DEGs were searched for homologues in *Arabidopsis* using Diamond (Buchfink *et al*., 2021), and genes of the best hit (the smallest E-value) were used for GO term enrichment analysis using g:Profiler (https://biit.cs.ut.ee/gprofiler/) with a threshold of a false discovery rate (FDR) < 0.05. Data sources of Gene Ontology (GO molecular function, GO cellular component, and GO biological process) and biological pathways (KEGG) were used. GO term sizes smaller than 10 and more than 500 were excluded from the analysis.

GO enrichment analysis results were visualized in Cytoscape (http://www.cytoscape.org/) application EnrichmentMap (http://www.baderlab.org/Software/EnrichmentMap).

We investigated the expression profiles of genes involved in (casparian strip) CS formation, including GRAS family transcription factor *SHR* and *SCR/SCARESCROW* (including *SCARECROW-LIKE* genes), MYB protein *MYB36*, receptor-like kinase *GSO1/SGN3/GASSHO1, CASPs*, protein kinase *SGN1*, peptides *CIF1/2*, peroxidase *PER64*, respiratory burst oxidase protein F *SGN4/RBOHF*, dirigent-like protein *ESB1*, and genes involved in suberin formation, including *β*-ketoacyl-CoA synthetase *KCS20*, fatty acyl ω-hydroxylase *CYP86* and *CYP86B1*, glycerol-3-phosphate acyltransferase *GAPT5*, dirigent-like protein *ESB1*, feruloyl transferase *ASFT*, GDSL-type esterase/lipase proteins, ATP-binding cassette *ABCG1/11/20*, lipid transfer protein *LTPI4* and *LTPG15*, MYB proteins *MYB39, MYB41, MYB70, MYB93, MYB107*, WRKY proteins *WRKY9, WRKY33*, and ANAC protein *NAC058*.

### Observation of CS and suberin deposition

Seed sterilization, germination, and hydroponic growth were the same as above. Salt treatment of 200 mM of NaCl started after two weeks. Roots in control and salt conditions were sampled at three, five, and seven days after salt treatment. Collected root samples were kept in 70 % ethanol at 4 ^o^C until sectioning. The main roots were sliced using a sharp double-edged razor blade under a stereomicroscope (Leica MZ16, Leica Microsystems) at 10 cm and 15 cm from the root apex. For observation of lignin deposition, root sections were stained by a mix of 3 % phloroglucinol and 35 % hydrochloric acid in a volume ratio of 2:1. For observation of suberin lamellae, root sections were incubated in a saturated solution of Sudan III at 70 °C for one minute, washed with distilled water for three times, and embedded in glycerol-water. The stained root sections were immediately observed under microscopy (Axioskop 2 plus, Zeiss).

### Piperonylic acid (PA) treatment

After two weeks of growth in hydroponics, plants were subjected to four conditions as follows: 1) Control (1 × Otsuka house); 2) Control + PA (1 × Otsuka house + 100 μM PA); 3) Salt (1 × Otsuka house + 200 mM NaCl); 4) Salt + PA (1 × Otsuka house + 200 mM NaCl + 100 μM PA). Treated plants were observed daily for any symptom of salt stress. Roots in all four conditions were sampled at three, five and seven days after treatments and kept in 70 % ethanol at 4 °C. After the treatments, Root sectioning, lignin and suberin deposition were observed as described above.

For determination of ions of plants, shoots and roots were collected seven days after the treatments, dried at 50 °C for one week, and then the weights were measured. Samples were digested with 69 % HNO_3_ for the determination of ions, including sodium (Na), magnesium (Mg), potassium (K), and Calcium (Ca), by inductively coupled plasma-mass spectrometry (ICP-MS, NexION 350S, PerkinElmer, Waltham, MA, USA).

## Results

### The vigor of the *Vigna* accessions under salt stress

To test the effect of 200 mM of NaCl on the vigor of the *Vigna* accessions, the symptoms of leaf wilting were evaluated in JP247202 (salt-tolerant accession of *V. marina*), JP233389 (salt-tolerant accession of *V. luteola*) and JP235855 (salt-sensitive accession of *V. luteola*). Consistent with our previous result of chlorophyll fluorescence change (Wang *et al*., 2023), the first leaf of JP235855 started wilting on day 1 and wilted on day 5 whereas no salt damage was detected for JP247202 and JP233389 until day 7 (Suppl. Fig. 1).

### Global feature of transcriptome data

To understand the mechanism of salt tolerance in JP247202, we analyzed the root transcriptome data of JP247202, JP233389 and JP235855 (Noda *et al*., 2024). Multidimensional scaling (MDS) plot revealed that both genotypic difference and salt treatment accounted for the variation in the transcriptome (Fig. 1a). The genotypic effect was apparent between JP247202 (*V. marina*) and the other two accessions (*V. luteola*). JP247202 formed distinct clusters from the other two accessions in both control and salt treatment, while JP233389 and JP235855 clustered closer to each other (Fig. 1a).

**Fig. 1.**
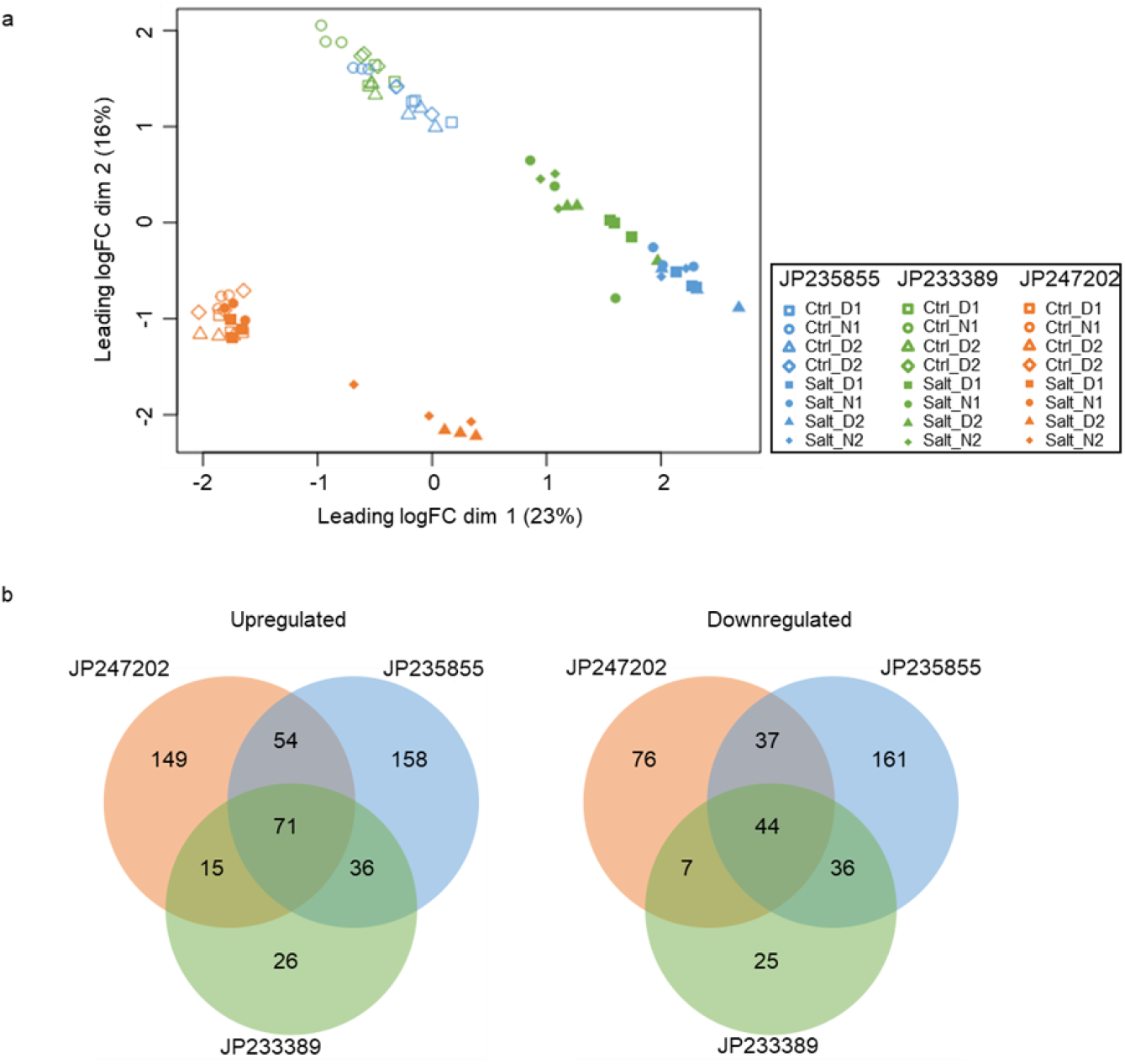
Multidimensional scaling (MDS) plot and common differentially expressed genes (DEGs) of JP235855, JP233389 and JP247202. **a**. MDS plot shows the similarities and differences between samples in genome-wide expression profiles, as indicated by die leading logFC (base 2 logarithm of fold change) dim (dimension) 1 and 2. Each point represents one sample and points with die same symbol represents three biological replicates. Blue, green, and orange indicates samples of JP235855, JP233389 and JP247202, respectively. Empty and filled symbols indicate samples in control (Ctrl) and salt treatment of 200 niM of NaCl (Salt), respectively. Square, circle, triangle and diamond indicate samples of the first day 1 (D1), first night (N1). second day (D2), and second night (N2). **b**. Venn diagram of upregulated and downregulated common DEGs in JP235855 (blue). JP233389 (green) and JP247202 (orange). Common DEGs are defined as those shared between time points (D1, N1, D2, and N2 for JP235855 and JP233389; D2 andN2 for JP247202).

For JP247202, the effect of salt treatment was not observed at the first day (D1) and first night (N1), as the transcriptomes in salt treatment clustered with those in control, but the effect was observed at the second day (D2) and second night (N2) (Fig. 1a). For JP233389 and JP235855, different from JP247202, the effect of salt treatment started from D1 (Fig. 1a).

### Differentially expressed genes (DEGs) analysis

Consistent with the global feature indicated by MDS, JP247202 at D1 had only one DEG between control and salt and no DEG at N1 (Suppl. Table 2). For downstream analyses, we focused on “common DEGs” that were shared between time points. Compared to the salt-sensitive JP235855, which had 597 common DEGs, the salt-tolerant JP247202 and JP233389 had fewer common DEGs, which were 453 and 260, respectively (Suppl. Table 2).

The overlap of common DEGs between accessions was shown in Fig. 1b. Among common DEGs in each accession, around 50% were JP247202- and JP235855-specific, whereas only ∼20% were JP233389-specific, indicating the transcriptomic response to salt in JP233389 was intermediate between that of JP247202 and JP235855.

### Gene Ontology (GO) enrichment analysis

Using common DEGs, GO enrichment analysis was carried out in JP247202, JP233389, and JP235855, for upregulated and downregulated genes separately. We identified GO terms of biological process (BP), molecular function (MF), cellular component (CC) and KEGG pathway that were specific to or common among accessions.

Consistent with the pattern of common DEGs (Fig. 1b), JP247202 and JP235855 had more accession-specific enriched GO terms than JP233389 (Suppl. Table 3). Only three terms were common between three accessions, including one BP term “secondary metabolic process” and two MF terms “carbohydrate transmembrane transporter activity” and “sugar transmembrane transporter activity” (Suppl. Table 3).

GO enrichment analysis highlighted the BP terms “phenylpropanoid metabolic process” and “phenylpropanoid biosynthetic process”, which were enriched in the two salt-tolerant accessions JP247202 and JP233389 (Suppl. Table 3).

For the KEGG pathway, two (“Cutin, suberine and wax biosynthesis” and “ABC transporters”) were specifically enriched in JP247202 and one (“Nitrogen metabolism”) was specifically enriched in JP235855 (Suppl. Table 3).

### Enrichment map for GO terms

Pathway information is redundant as enrichment analysis often results in several versions of the same pathway (Reimand *et al*., 2019). To simplify interpretation, enrichment maps were used to collapse redundant BP and MF terms, respectively, into single biological themes.

Enriched BP terms formed five networks (Fig. 2). The first network included six terms (Fig. 2). The first four terms related to metabolic and biosynthetic processes of secondary metabolite including phenylpropanoid, were shared by the salt-tolerant JP247202 and JP233389, indicating these two accessions have some common mechanisms for salt tolerance. The latter two terms, “suberin biosynthetic process” and “lignin metabolic process”, were JP247202-specific (Fig. 2).

**Fig. 2.**
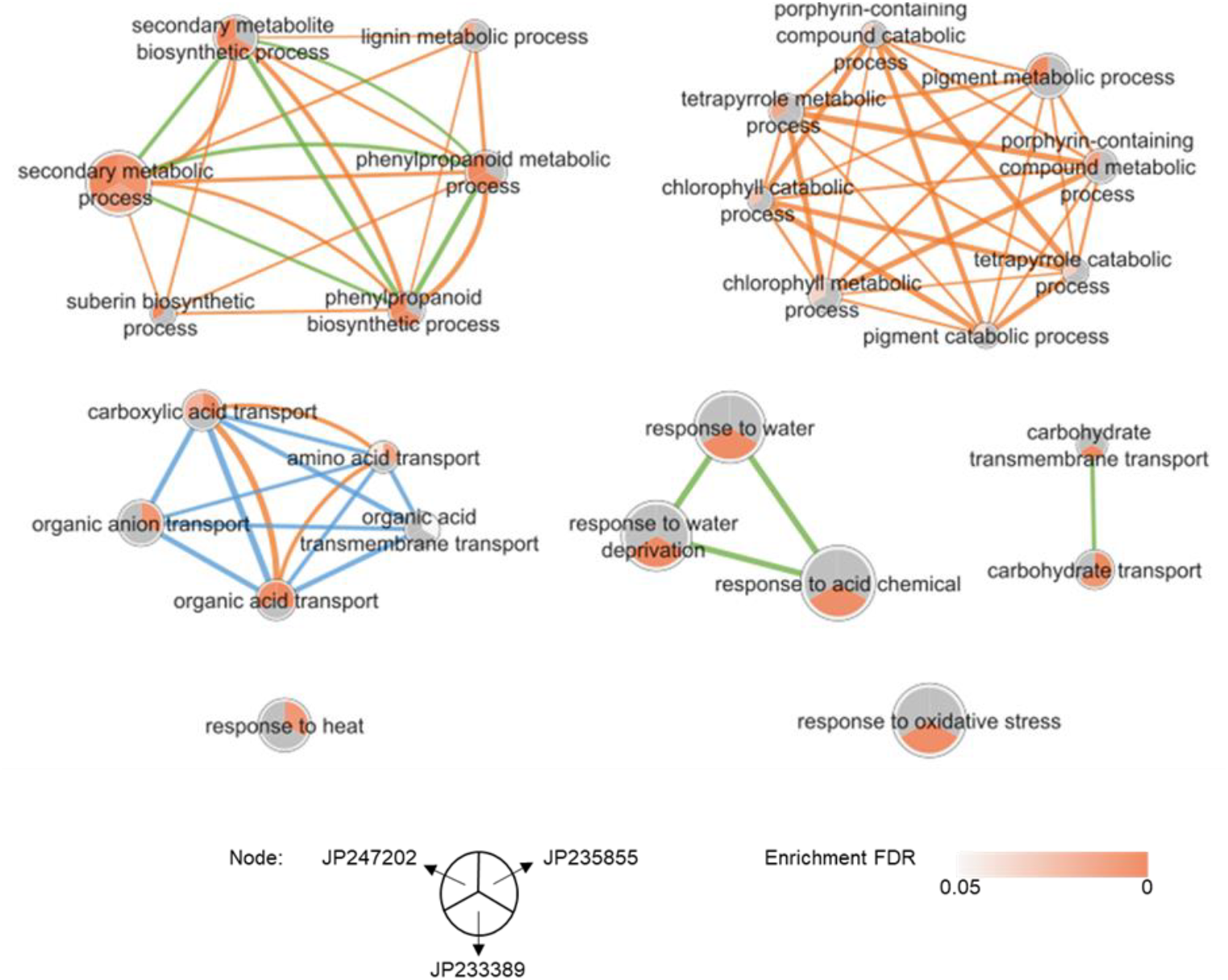
Enrichment map for biological process (BP) terms enriched in JP235855, JP233389 and JP247202. BP terms are shown as nodes that are connected with edges if two BP terms share genes. Nodes are divided into three parts for JP235855, JP233389 and JP247202, as illustrated at the bottom of the figure. Nodes are sized by the number of genes and colored by false discovery rate (FDR) of enrichment (grey color indicate no enrichment in specific accession). Edges are colored for JP235855 (blue), JP233389 (green) and JP247202 (orange), respectively.

The second network was JP247202-specific, with eight terms related to metabolic and catabolic processes of pigment, including porphyrin-containing compound, tetrapyrrole, and chlorophyll (Fig. 2).

The third network included five terms involving transport of organic acid, carboxylic acid, amino acid, organic anion, and organic acid transmembrane transport. The first three were shared by the salt-tolerant JP247202 and salt-sensitive JP235855, and the latter two terms were JP235855-specific (Fig. 2).

The fourth and fifth networks were JP233389-specific, including terms of response to water and acid chemical and carbohydrate transport, respectively (Fig. 2).

Enriched MF terms formed four networks. The first network was JP247202-specific including five terms related to activity of phosphatidylinositol-phosphate-phosphatase (Suppl. Fig. 2).

The second network included five terms related to transmembrane transporter activity of carbohydrate, sugar, sucrose, oligosaccharide, and disaccharide. While the first two were shared by three accessions, the later three terms were JP233389-specific (Suppl. Fig. 2)

The third and fourth networks were JP235855-specific, including terms related to activity of dioxygenase, 2-oxoglutarate-dependent dioxygenase, oxidoreductase and monooxygenase, and terms related to transmembrane transporter activity of amino acid, carboxylic acid and organic acid, respectively (Suppl. Fig. 2).

### Promising network and likely candidate genes

The first network of BP terms led us to a hypothesis on the formation of root barriers. The phenylpropanoid pathway generates the building blocks for lignin and suberin (Cesarino *et al*., 2022). Lignin, in the form of CS, and suberin lamellae are apoplastic barriers to the flow of ion. Therefore, CS and suberin may function as barriers to Na^+^ loading to the xylem, especially in JP247202.

We investigated genes that enriched in the first network of BP terms (Fig. 2), and five genes were pulled out as they had higher expression levels in the salt-tolerant JP247202 than those in the other two accessions (Fig. 3a, Suppl. Table 4). Two genes were involved in the biosynthetic process of lignin: *OMT1*, a O-methyltransferase, and *GT72B1*, a glycosyltransferase. We also found *ABF4* that encodes a bZIP transcription factor and is involved in the ABA-activated signaling pathway.

**Fig. 3.**
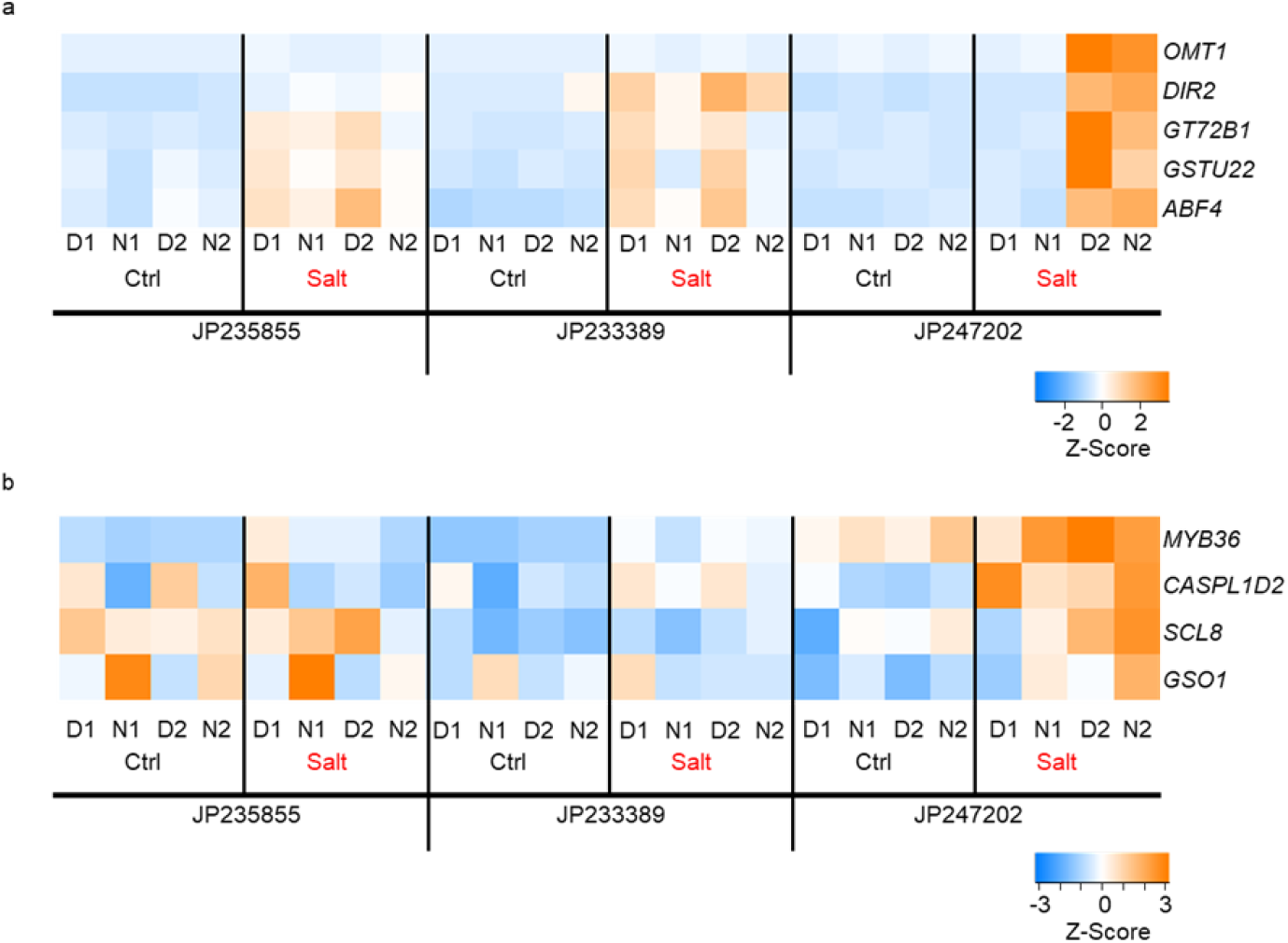
Heatmap of gene expressions in JP235855, JP233389 and JP247202 under control (Ctrl) and 200 mM of NaCl (Salt). **a**. Genes enriched the BP terms of the first network and with higher expression in JP247202. **b**. Gene involved in the formation of casparian strip (CS). Heatmap is colored by Z-Score normalized trimmed mean of M values (TMM).

In case we miss genes that have not been assigned with BP terms in the promising network, we directly investigated homologous genes involved in CS formation (Suppl. Table 4, and see Materials and Methods). As a result, *MYB36, CASPL1D2, SCL8*, and *GSO1*, had higher expression levels in JP247202 than the other two accessions or were salt induced only in JP247202 (Fig. 3b).

We also investigated homologous genes involved in suberin formation (see Materials and Methods) but none of them were differentially expressed among the accessions and responded to salt stress.

### Observations on root barriers

As the transcriptome data suggested salt-induced formation of CS and suberin lamella in JP247202, we examined the presence of lignified CS and suberin deposition at root endodermis. As a result, although CS formation on day 5 and 7 was salt induced at 10 cm from root apex in all three accessions, differences were obvious between accessions (Fig. 4a). The salt-sensitive JP235855 formed only spot-like CS on a few endodermal cells whereas the salt-tolerant JP233389 and JP247202 formed CS in an entire circle in the endodermis (Fig. 4a). Interestingly, while JP233389 formed the typical structure of Casparian bands, JP247202 expanded the deposition of lignin to the outer- or inner-side of the endodermal cells (Fig. 4a). In addition, the endodermis-like cells were partially bi-layered (Fig. 4a). This trend was even more obvious in the 15 cm position from the root apex, where JP247202 deposited lignin in ∼3 layers of cells around endodermis (Suppl. Fig. 3), while JP233389 and JP235855 deposited in one-layered or partially bi-layered cells of endodermis. Thus, in response to salt stress, JP247202 built up thick and dense apoplastic barrier around endodermis.

**Fig. 4.**
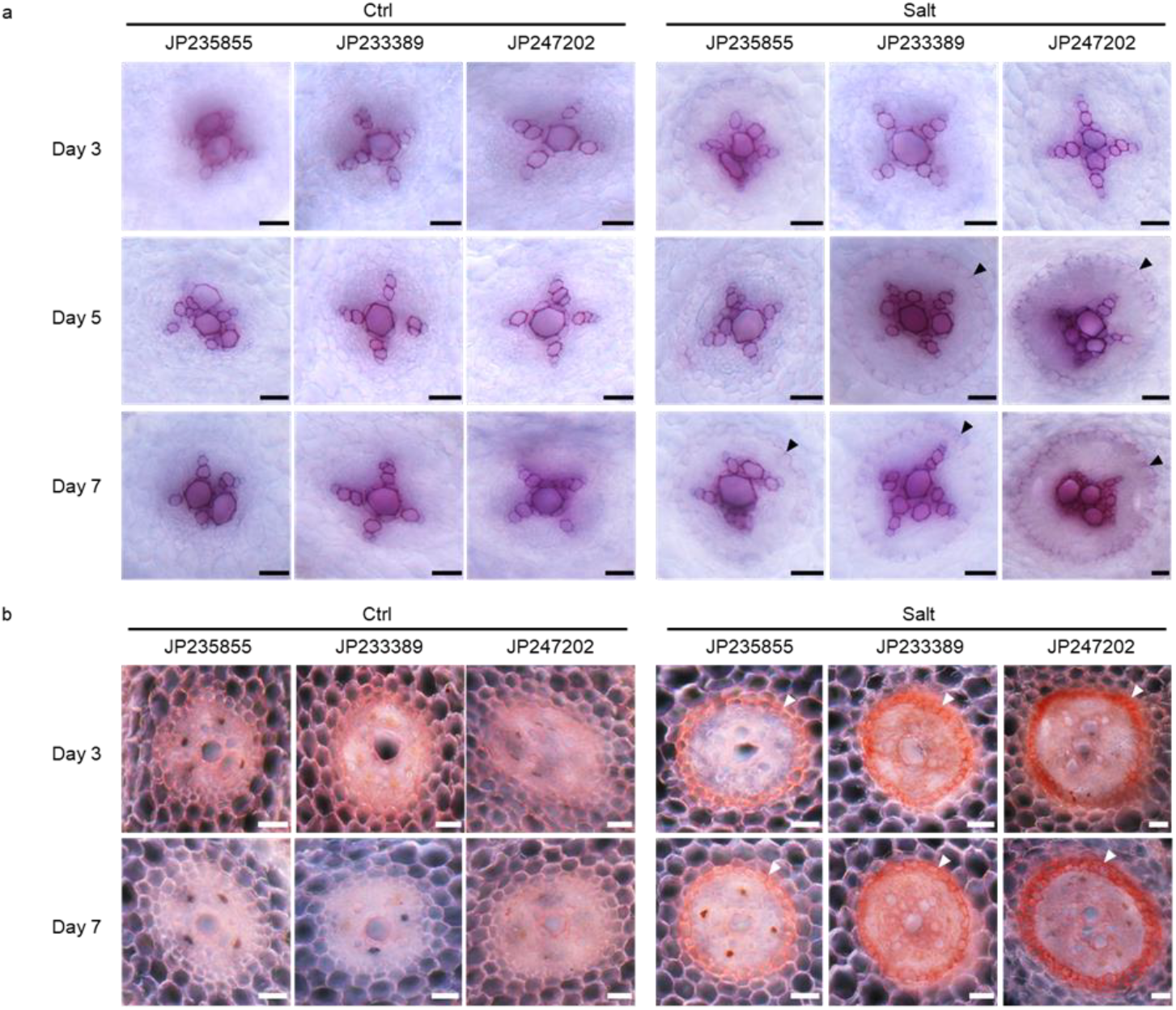
Staining of root barriers in JP235855, JP233389 and JP247202 under control (Ctrl) and 200 inM of NaCl (Salt). **a**. Staining of lignin, **b**. Staining of suberin lamella. Roots were collected at three, five and seven days of salt treatment. Roots were sectioned at 10 cm from apex. Scale bar = 50 μm. Black and white arrow-heads indicate CS and suberin deposition, respectively.

The formation of suberin lamella was detected three days after the initiation of salt stress (Fig. 4b). Compared to the salt-tolerant JP247202 and JP233389, deposition of suberin was much weaker in the salt-sensitive JP235855.

### Effect of apoplastic barrier on Na^+^ transport

To test the role of apoplastic barrier in Na^+^ loading, we tried to disrupt the formation of lignin by exogenous treatment with piperonylic acids (PA), a C4H-inhibitor that inhibits monolignol formation and thus inhibits lignification of CS (Naseer *et al*., 2012; Lee *et al*., 2013; Duan *et al*., 2018; Andersen *et al*., 2021). We confirmed that PA inhibited salt-induced formation of lignin in JP233389 and JP247202, but did not affect suberin deposition (Fig. 5a, b, Suppl. Fig. 4a, b).

**Fig. 5.**
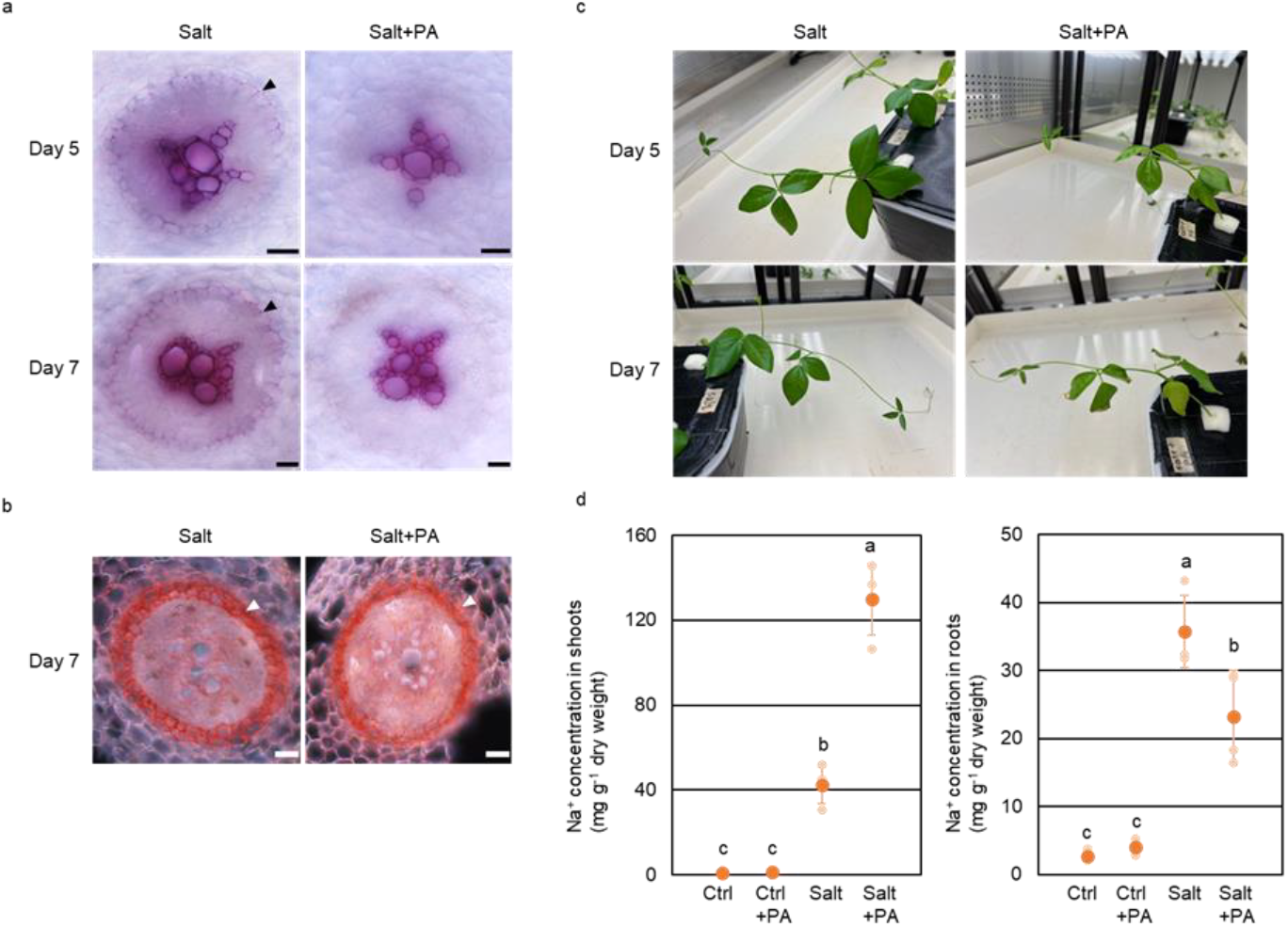
Effect of defective apoplastic barrier on plant growth and Na^+^ allocation in JP247202. **a**. Staining of lignin in JP247202 under 200 mM of NaCl (Salt) and Salt + PA (100 μM). **b**. Staining of suberin lamella in JP247202 under 200 mM of Salt and Salt + PA (100 μM). PA: Piperonylic acid. Roots were collected at five and seven days of treatment. Roots were sectioned at 10 cm from apex. Scale bar = 50 μm. Black and white arrow-heads indicate lignin and suberin deposition, respectively, **c**. Photos of JP247202 at five and seven days of treatments. Leaves in Salt + PA show symptoms of salt damage, d. Na^+^ concentration in shoots and roots of JP247202 at seven days of treatments. Treatments include control (Ctrl). Ctrl + PA. Salt and Salt + PA. Means are calculated using four replications and those not sharing the same alphabet are statistically different (Tukey LSD test. P < 0.05).

We observed symptoms of salt damage, including etiolated, winkled, and burning leaves, in the salt-tolerant JP247202 with the “Salt + PA” treatment, whereas plants with only salt treatment did not show such symptoms (Fig. 5c).

Supporting our hypothesis, plants treated with PA dramatically increased Na^+^ allocation to shoots (Fig. 5d, Suppl. Fig. 4c). Shoots of JP247202 in “Salt + PA” had more than three times higher Na^+^ (130 mg g^-1^) than that in “Salt” (Fig. 5d). Similarly, another salt-tolerant accession JP233389 in “Salt + PA” had 74 mg Na^+^ per dry weight shoots, which was 2.6 times higher than that in “Salt” (Suppl. Fig. 4c). Roots in “Salt + PA” had similar (JP233389) or lower (JP247202) Na^+^ concentration compared to that in “Salt” (Fig. 5d, Suppl. Fig. 4c).

### Nutrient transport under salt stress

Given that the reinforced lignin deposition was important in suppressing Na^+^ allocation to shoots, we were intrigued to ask whether it also affects the transport of other nutrients. Thus, we assessed K^+^, Mg^2+^, and Ca^2+^ in the roots and the shoots of JP233389 and JP247202.

As a result, K^+^ concentration in roots was significantly lower in JP247202 than in JP233389 under the control condition, but not different under the salt-stressed condition (Fig. 6a). Moreover, K^+^ concentration in shoots was not significantly different under the control condition but significantly higher in JP247202 than in JP233389 under the salt-stressed condition (Fig. 6b). The K^+^ concentration in roots also significantly decreased in response to salt stress in both accessions (Fig. 6a), while that in shoots did not significantly change (Fig. 6b). In addition, PA treatment further decreased the K^+^ concentration in roots under salt stress, although the decrease was not significant for JP247202 (Fig. 6a), while it did not significantly affect K^+^ concentration in shoots (Fig. 6b).

**Fig. 6.**
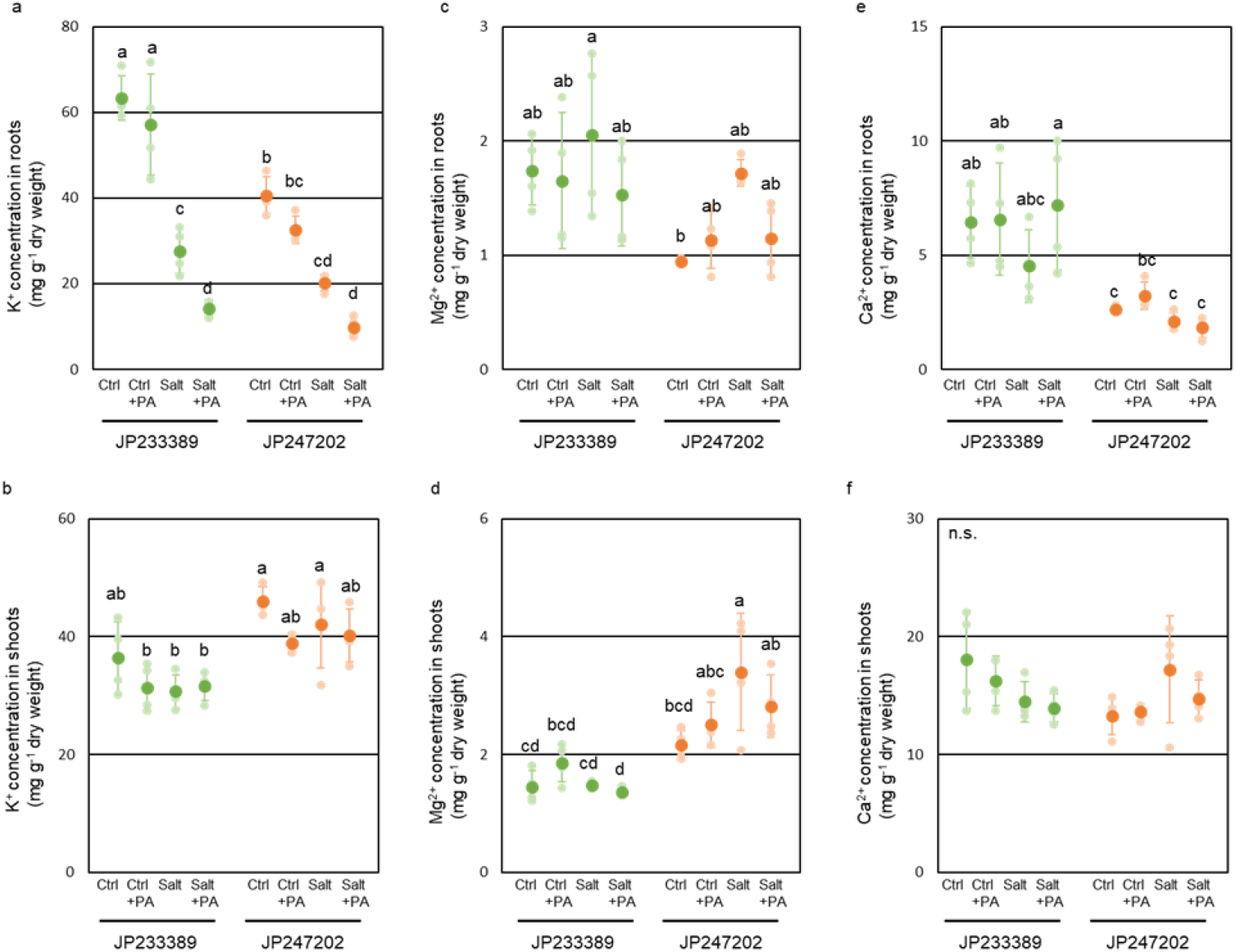
Ion concentrations in JP233389 and JP247202 under four conditions including control (Ctrl), Ctrl + PA (100 μM), 200 mM of NaCl (Salt), and Salt + PA (100 μM). **(a, b)** K^+^ concentration in roots (a) and shoots (b). **(c, d)** Mg^2+^ concentration in roots (c) and shoots (d). **(e, f)** Ca^2+^ concentration hi roots (e) and shoots (f). Roots and shoots were collected at seven days after treatments. PA: Piperonylic acid. Means are calculated using four replications and those not sharing the same alphabet are statistically different (Tukey LSD test, P < 0.05).

Mg^2+^ concentration in roots was not significantly different between JP247202 and JP233389 under both the control and salt treatment (Fig. 6c). Mg^2+^ concentration in shoots was also not different under the control condition, but was significantly higher in JP247202 than in JP233389 under the salt-stressed condition (Fig. 6d). In response to salt stress, Mg^2+^ concentration significantly increased in shoots of JP247202 but not of JP233389 (Fig. 6d). Additionally, PA treatment did not affect Mg^2+^ concentration in roots and shoots of both accessions (Fig. 6c, d).

Ca^2+^ concentration was significantly lower in roots of JP247202 in both control and salt treatment (Fig. 6e), while it was not significantly different across accessions or treatments in shoots (Fig. 6f).

As JP247202 maintained nutrient transport even with thicker apoplastic barrier, we looked into the transcriptome data and checked the transcription profiles of the genes involved in nutrient transport. As a result, we found, in response to salt stress, JP247202 up-regulated potassium transporters KUP6 and KT3, positive regulator of the potassium transporter CIPK23, calcium transporter interacting protein GRX2, and magnesium transporters MGT10, AVI2 (Suppl. Fig. 5).

## Discussion

In this study, we elucidated the importance of apoplastic barrier in the excellent ability of Na^+^ exclusion in JP247202 of *V. marina*, an accession of the most salt-tolerant species in the genus *Vigna*. When salt-stressed, JP247202 deposited lignin in multi-layered cells around endodermis, thereby effectively suppressing Na^+^ transport from roots to shoots. We also found that the reinforced apoplastic barrier did not lead to a reduction in the accumulation of K^+^, Mg^2+^, and Ca^2+^ in the shoots of JP247202 under salt stress.

### Salt-induced lignin formation is a primary barrier to the transport of Na^+^

Our results strongly indicated the apoplastic barrier is essential for restricting Na^+^ transport. Studies on *Arabidopsis* mutants and Maize inbred lines have revealed that CS plays vital roles in restricting transpiration-dependent sodium uptake (Reyt *et al*., 2021; Calvo‐Polanco *et al*., 2021; Wang *et al*., 2022). Our results further highlighted the role of lignified apoplastic barrier in salt tolerance, as JP247202 and JP233389 almost totally lost salt tolerance with defective lignification (Fig. 5, Suppl. Fig. 4). Also, the thickness of lignin barrier was correlated to the ability of suppressing Na^+^ transport. As shown in Fig. 4, JP247202 formed multi-layered apoplastic barrier, whereas JP233389, which is relatively less salt-tolerant, formed typical, one-layered Casparian bands, and the salt-sensitive JP235855 formed only spot-like CS. The reinforced apoplastic barrier and high ability of suppression of Na^+^ transport enables JP247202 to dominate tropical marine beaches, which is not only the environment of high salinity but that of high transpiration.

However, we were first puzzled when we observed “something more than CS” in the endodermis of JP247202. As described in the existing literatures (Miyashima and Nakajima, 2011; Geldner, 2013), we have believed that seed plants typically have endodermis as single layer and CS forms in between the endodermal cells. However, in JP247202, lignin is deposited in all sides of the endodermal cells. Moreover, the cell wall lignification expanded to several layers of cells, though currently we do not know whether it is multi-layered endodermis or ectopic lignification in cortical cells. In any case, the expanded lignification around endodermis of JP247202 should greatly reinforce the apoplastic barrier to any soluble ions including Na^+^.

The reinforced apoplastic barrier in JP247202 under salt stress corresponded well with the expression profiles of lignin biosynthetic genes (Fig. 3). Such genes include *Myb36*, the master regulator of CS formation (Kamiya *et al*., 2015), and *CASPL1D2* (*CASP-LIKE PROTEIN 1D2*), a CASPs protein family involved in initiating CS formation (Roppolo *et al*., 2011). Moreover, *SCARECROW-like 8* (*SCL8*), which determines cell fate of endodermal cells (Nakajima *et al*., 2001), was strongly induced by salt stress in JP247202 (Fig. 3b), possibly leading to proliferation of endodermis-like cells and lignin deposition in additional layers of cells. Additionally, *GSO1*, involved in CS assembly and endodermal cell differentiation (Pfister *et al*., 2014), was induced by salt stress only in JP247202 (Fig. 3b), further reinforcing formation of apoplastic barrier.

While we revealed that the reinforced apopalstic barrier is essential for suppression of Na^+^, we do not conclude it explains all aspects of sodium exclusion in JP247202. In the apical part, where root cells undergo cell division and elongation, the apoplastic barrier should not form (Geldner, 2013). Thus, the apoplastic transport of Na^+^ cannot be suppressed by CS even in the salt-tolerant JP247202. There, Na^+^ excretion by SOS1 (Noda *et al*., 2024) could be crucial for the restriction of Na^+^ accumulation. Moreover, the excess amount of Na^+^ that enters into the cortical cells is required to be extruded out of the cells to maintain ion homeostasis and nutrient uptake capacity (Essa, 2002). The reinforced apoplastic barrier and high ability of Na^+^ excretion may synergistically enforce the salt tolerance of JP247202.

### Salt-induced apoplastic barrier does not prevent the transport of K^+^, Ca^2+^, Mg^2+^

When we observed the thick lignified barrier formed in JP247202 under salt stress, we wondered that it could restrict the transport of not only Na^+^ but other nutrients. However, even with thicker lignified barrier compared to JP233389 (Fig. 4), JP247202 under salt stress transported higher amount of K^+^ and Mg^2+^ to shoots (Fig. 6b, d). Similarly, despite thicker lignified barrier in the salt stress than in the control condition (Fig. 4), the nutrient transported to shoots remained unchanged in JP233389 and even increased in JP247202 (Fig. 6b, d, f). Interestingly, the transport of the essential ions to shoots was not affected by defect of the apoplastic barrier (Fig. 6b, d, f). These results indicate that symplastic pathway should play major role in transporting such essential nutrients and its capacity is maintained or even enhanced by salt stress in JP247202. The up-regulation of nutrient transporters (Suppl. Fig. 5) further supports this idea, though post-translational regulation may also contribute to nutrient uptake.

### Salt-induced suberin deposition is not the key to suppress the transport of Na^+^

We also found the formation of suberin lamella was not enough to restrict Na^+^ transport, as PA treatment, which led to significant increase of Na^+^ accumulation in shoots, disturbed only deposition of lignin and not suberin deposition in endodermal cells (Fig. 5, Suppl. Fig. 4). It has long been argued that suberin lamella is also important for controlling Na^+^ transport across endodermis (Wang P. *et al*., 2020; Shukla *et al*., 2021), but our results support the recent study by Calvo-Polanco *et al*. (2021) that CS is more important for restricting Na^+^ transport.

Instead, we speculate that the salt-induced formation of suberin lamella in JP247202 may contribute to the ability to retain water uptake (Yoshida *et al*., 2020, Wang *et al*., 2023), rather than to ion transport. However, currently we do not have evidence for it and need further study to test this hypothesis.

### Other JP247202-specific GO network

JP247202 under salt stress was specifically enriched for the activity of “phosphatidylinositol phosphate phosphatase” (Suppl. Fig. 2). The phosphatidylinositol-4,5-biphosphate controls membrane trafficking (Jost *et al*., 1998; Li *et al*., 2020) and phosphatidylinositol-3,4-biphosphate controls the spatiotemporal endocytosis (Posor *et al*., 2013). As endomembrane trafficking is important in plants in response to salt stress (Wang X. *et al*., 2020), our result suggested a putative role of phosphatidylinositol phosphate phosphatase in salt tolerance of JP247202.

Another JP247202-specific GO network related to metabolic and catabolic processes of chlorophyll (Fig. 2), however, might be over-representative for the root transcriptome, though chlorophyll fluorescence is an indicator of photosynthetic activity and a signal of stress response (Guidi *et al*., 2019).

In conclusion, our transcriptome and histological analyses strongly indicate that multi-layered apoplastic barrier is critical for sodium exclusion of *V. marina*. Our results also revealed that lignin deposition is more important than suberin lamellae for restricting Na^+^ transport. In addition, we found *V. marina* is able to maintain transport of essential ions even under salt stress, presumably via enhanced symplastic transport. Thus, to develop a salt-tolerant crop, it might be essential to have a barrier for sodium and paths for nutrients.

## Acknowledgements

We appreciate Dr. Jimmy Burridge for his kind advice and instruction on root sectioning.

## Author Contributions

Wang F. and Naito K. planned and designed the research. Wang F. performed transcriptome analyses. Wang F. and Yamauchi T. performed root sectioning and staining. Tanoi K. performed ion assessment. Wang F. wrote the original manuscript. Naito K., Yamauchi T. and Tanoi K. reviewed and edited the manuscript.

