## Supplemental Figures for "Multi-layered apoplastic barrier underlying the ability of Na^+^ exclusion in *Vigna marina*"

### Slide 1
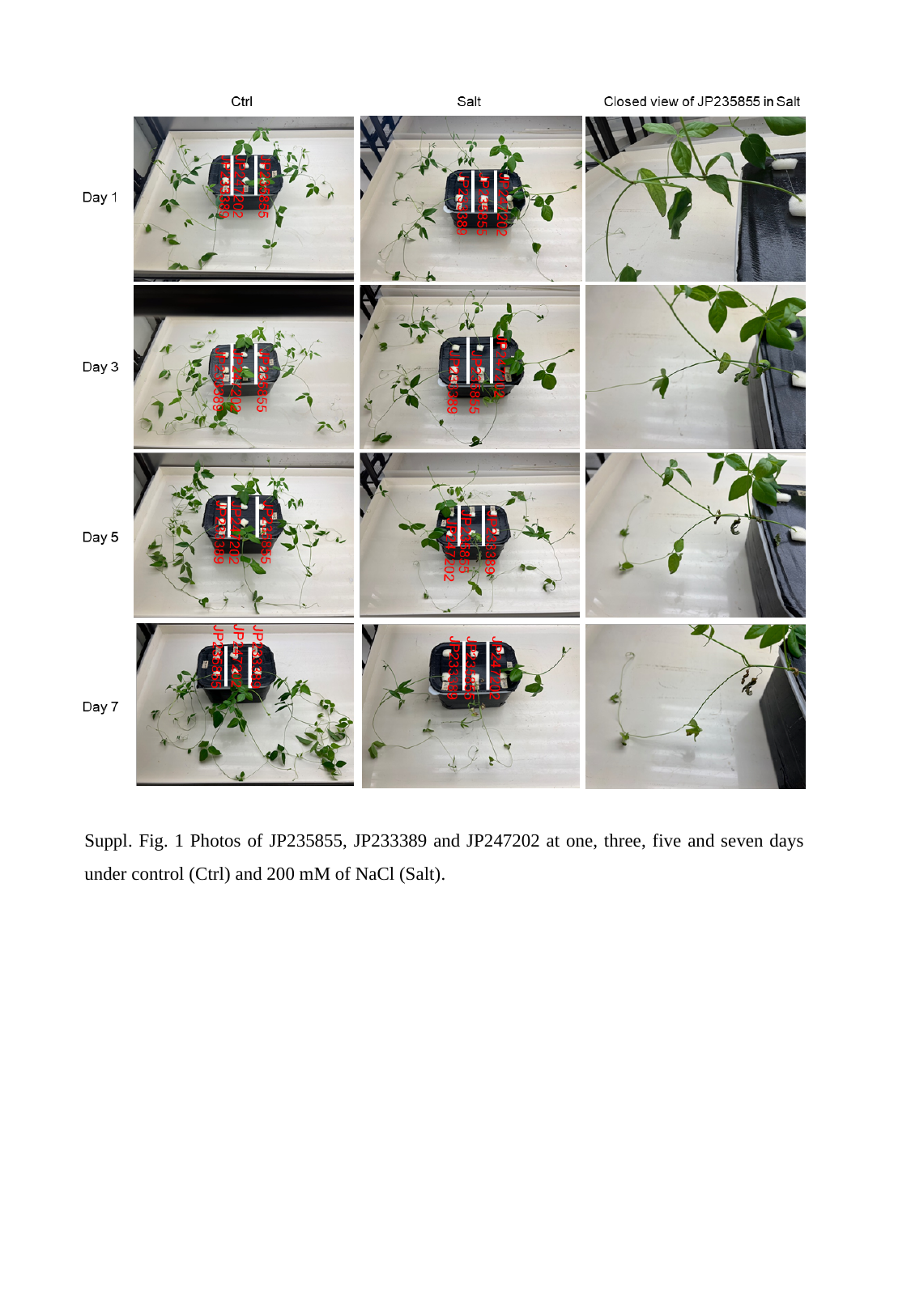

Suppl. Fig. 1 Photos of JP235855, JP233389 and JP247202 at one, three, five and seven days under control (Ctrl) and 200 mM of NaCl (Salt).

### Slide 2
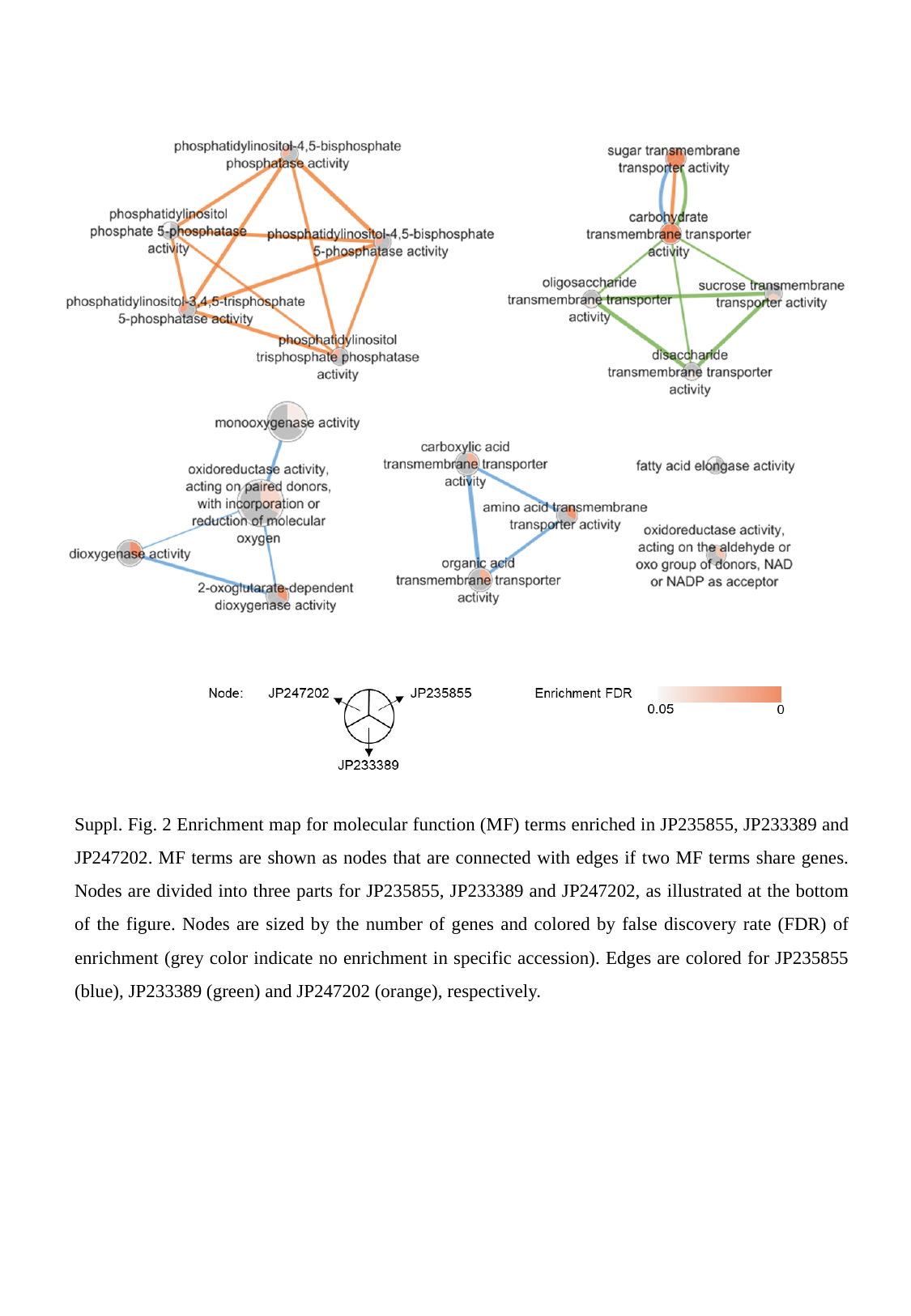

Suppl. Fig. 2 Enrichment map for molecular function (MF) terms enriched in JP235855, JP233389 and JP247202. MF terms are shown as nodes that are connected with edges if two MF terms share genes. Nodes are divided into three parts for JP235855, JP233389 and JP247202, as illustrated at the bottom of the figure. Nodes are sized by the number of genes and colored by false discovery rate (FDR) of enrichment (grey color indicate no enrichment in specific accession). Edges are colored for JP235855 (blue), JP233389 (green) and JP247202 (orange), respectively.

### Slide 3
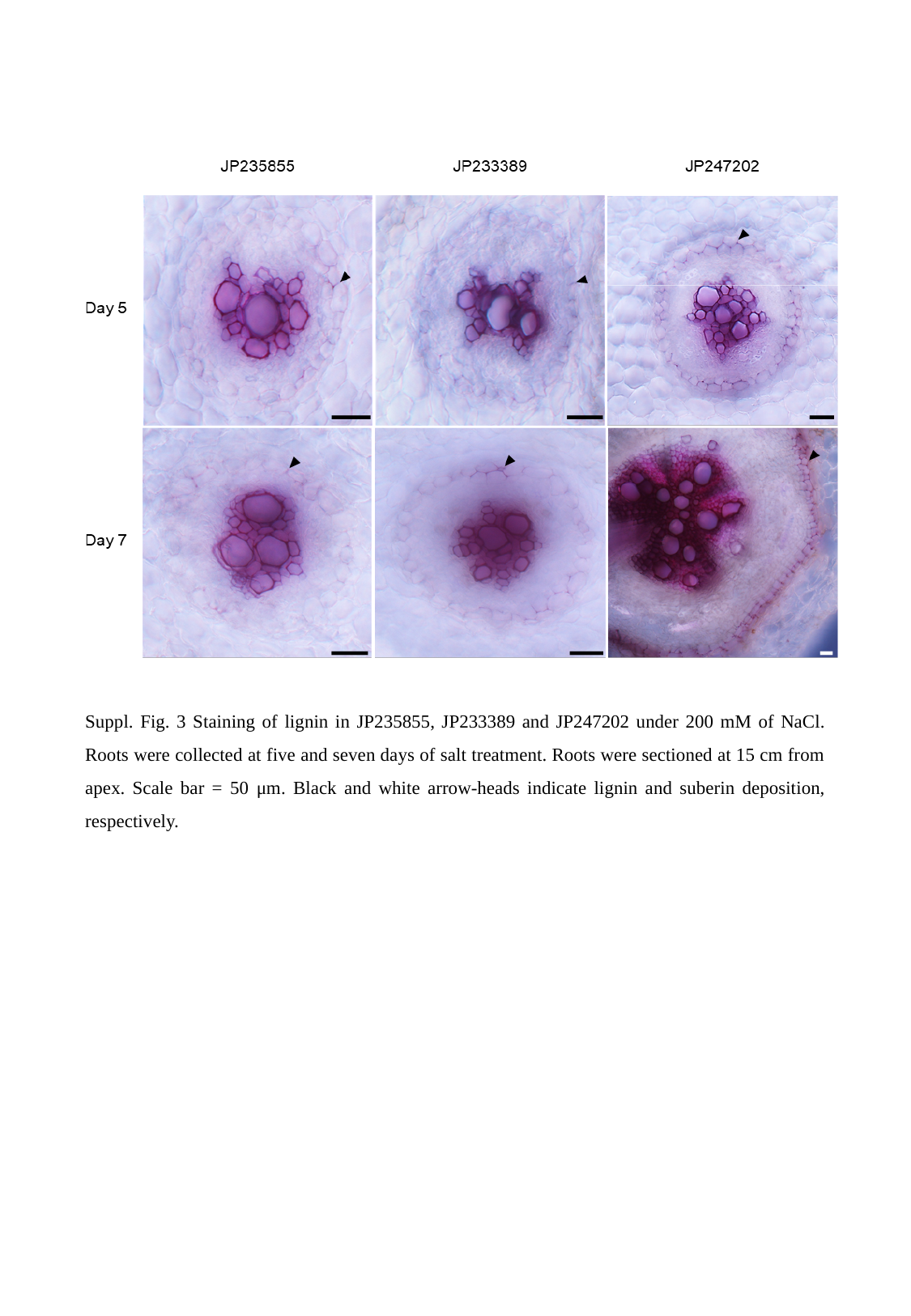

Suppl. Fig. 3 Staining of lignin in JP235855, JP233389 and JP247202 under 200 mM of NaCl. Roots were collected at five and seven days of salt treatment. Roots were sectioned at 15 cm from apex. Scale bar = 50 μm. Black and white arrow-heads indicate lignin and suberin deposition, respectively.

### Slide 4
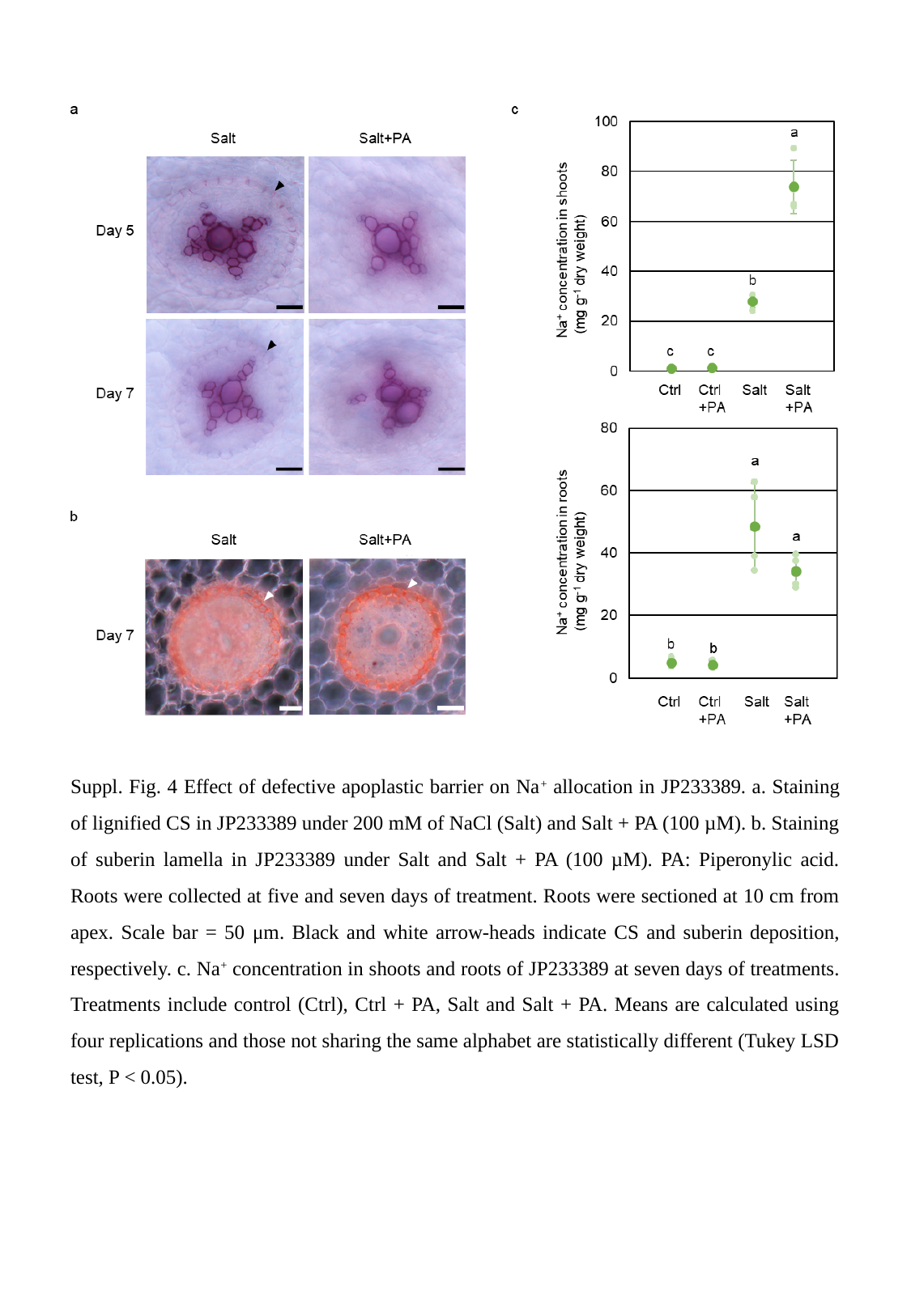

Suppl. Fig. 4 Effect of defective apoplastic barrier on Na^+^ allocation in JP233389. a. Staining of lignified CS in JP233389 under 200 mM of NaCl (Salt) and Salt + PA (100 µM). b. Staining of suberin lamella in JP233389 under Salt and Salt + PA (100 µM). PA: Piperonylic acid. Roots were collected at five and seven days of treatment. Roots were sectioned at 10 cm from apex. Scale bar = 50 μm. Black and white arrow-heads indicate CS and suberin deposition, respectively. c. Na^+^ concentration in shoots and roots of JP233389 at seven days of treatments. Treatments include control (Ctrl), Ctrl + PA, Salt and Salt + PA. Means are calculated using four replications and those not sharing the same alphabet are statistically different (Tukey LSD test, P < 0.05).

### Slide 5
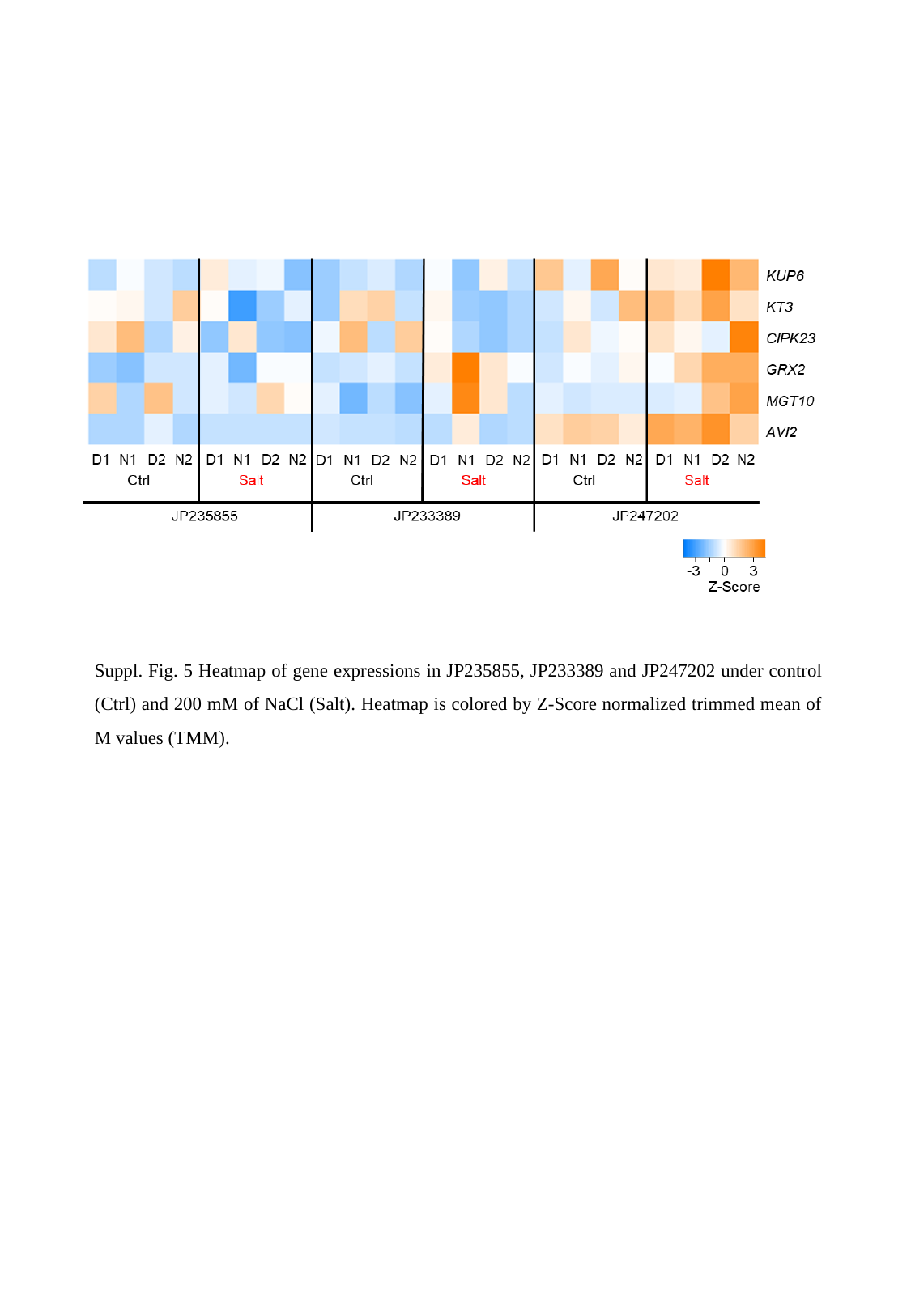

Suppl. Fig. 5 Heatmap of gene expressions in JP235855, JP233389 and JP247202 under control (Ctrl) and 200 mM of NaCl (Salt). Heatmap is colored by Z-Score normalized trimmed mean of M values (TMM).
